# Structural basis of Irgb6 inactivation by *Toxoplasma gondii* through the phosphorylation of switch I

**DOI:** 10.1101/2022.10.31.514472

**Authors:** Hiromichi Okuma, Yumiko Saijo-Hamano, Aalaa Alrahman Sherif, Emi Hashizaki, Naoki Sakai, Takaaki Kato, Tsuyoshi Imasaki, Eriko Nitta, Miwa Sasai, Yoshimasa Maniwa, Hidetaka Kosako, Daron M Standley, Masahiro Yamamoto, Ryo Nitta

## Abstract

Upon infection with *Toxoplasma gondii*, host cells produce immune-related GTPases (IRGs) to kill the parasite. *T. gondii* counters this response by releasing ROP18 kinase, which inactivates IRG GTPases and inhibits their recruitment to the *T. gondii* parasitophorous vacuole (PV). However, the molecular mechanisms of this process are entirely unknown. Here we report the atomic structures of Irgb6 with a phosphomimetic mutation by ROP18. The mutant has lower GTPase activity and is not recruited to the PV membrane (PVM). The crystal structure shows the mutant exhibit a distinct conformation from the physiological nucleotide-free form, thus preventing GTPase cycling. This change allosterically modifies the conformation of the membrane-binding interface, preventing physiological PVM-binding. Docking simulation of PI5P also supports the impaired binding of the mutant to PVM. We thus demonstrate the structural basis for *T. gondii* escape from host cell-autonomous defense, and provide a structural model for regulating enzymatic activity by phosphorylation.

## Introduction

*Toxoplasma gondii* is an important pathogen for warm-blooded animals including humans. It enters host cells and forms a membranous structure called parasitophorous vacuole (PV) in order to resist host defenses [1–2]. In response to *T. gondii* infection, host immunity releases IFN-γ, which cell-autonomously suppresses the intracellular growth of *T. gondii* and eventually kill the microorganism [3–5]. One prominent IFN-γ inducible event is the upregulation of two subfamilies of IFN-inducible GTPases, immunity-related GTPases (IRGs), and guanylate-binding proteins (GBPs). They specifically target the PV membrane (PVM) and disrupt the PV in which *T. gondii* is enfolded [6–10]. Among IFN-inducible GTPases, Irgb6 directly recognizes PI5P, which is accumulated on *T. gondii* PVM, and plays a pioneering role in recruiting other IFN-inducible GTPases such as Irga6, Irgb10 and GBPs and effectors such as p62/Sqstm1 and ubiquitin to the PVM [11–12).

In order to counter the host cell-autonomous defense system, *T. gondii* secretes various effector proteins including a serine-threonine kinase, ROP18 [13, 14]. The parasite-derived ROP18 phosphorylates host IRGs including Irgb6 at two conserved threonine residues in the switch I region, biochemically abrogating the GTPase activity of IRGs and inhibiting the recruitment of IRGs, thus rendering IRGs inactive. Although the inactivation of IRGs by ROP18 is well established by biochemical and cell biological evidence, the molecular mechanisms by which phosphorylation of the switch I region inhibits the GTPase activity of IRGs, or why such phosphorylated IRGs are no longer recruited to the PV membrane, are entirely unknown.

To resolve the structural mechanisms of IRG inactivation by parasite-derived ROP18, we solved the crystal structures of Irgb6 with a phosphomimetic mutation. As expected, the mutant was no longer recruited to the PVM and had inactivated GTPase activity. Moreover, the crystal structure showed that the G-domain of the mutant did not exhibit the nucleotide-free form, thus preventing GTPase cycling. More surprisingly, the conformational change of the G-domain allosterically modified the conformation of the membrane-binding interface on the opposite side of the protein, preventing physiological PI5P binding.

## Results

### Phosphorylation of Thr95 in switch I inhibits Irgb6 recruitment to *T. gondii* PVM

To assess the effect of phosphorylation of the Thr95 on Irgb6, the amino acid was substituted to aspartate as a phosphomimetic mutation (T95D mutation), following a previous paper [13]. We confirmed that phosphorylation of Thr95 was induced by *T. gondii* infection in MEF cells (S1 Fig). The Flag-tagged wild-type or T95D mutant of Irgb6 (Irgb6-T95D) was expressed in Irgb6-deficient MEFs (Fig 1A). An indirect immunofluorescence study showed that Flag-tagged wild-type Irgb6 (Irgb6-WT) was loaded onto *T. gondii* PVM (Fig 1A). In sharp contrast, Flag-tagged Irgb6-T95D was not detected on *T. gondii* PVM at all (Fig 1A), suggesting that the phosphorylation of Thr95 of Irgb6 is required for the localization on the PVM. Next, we examined the effect of Thr95 phosphorylation on recruitment of other IFN-inducible GTPases (Fig 1B). The recruitment of Irga6 and Irgb10 was not observed upon reconstitution of Irgb6-T95D in Irgb6-deficient MEFs (Fig 1B). IFN-γ stimulates coating of effectors such as ubiquitin and p62/Sqstm1 on *T. gondii* PVM in a manner dependent on IFN-inducible GTPases [15–16]. Therefore, we next analyzed whether loading of ubiquitin and p62/Sqstm1 is affected by the Thr95 phosphorylation (Fig 1C). Although reconstitution of Irgb6-WT recovered effector loading on *T. gondii* PVMs in Irgb6-deficient MEFs, reconstitution of Irgb6-T95D did not (Fig 1C). Taken together, these data indicate that Thr95 phosphorylation of Irgb6 negates recruitment to *T. gondii* PVM.

**Fig 1.**
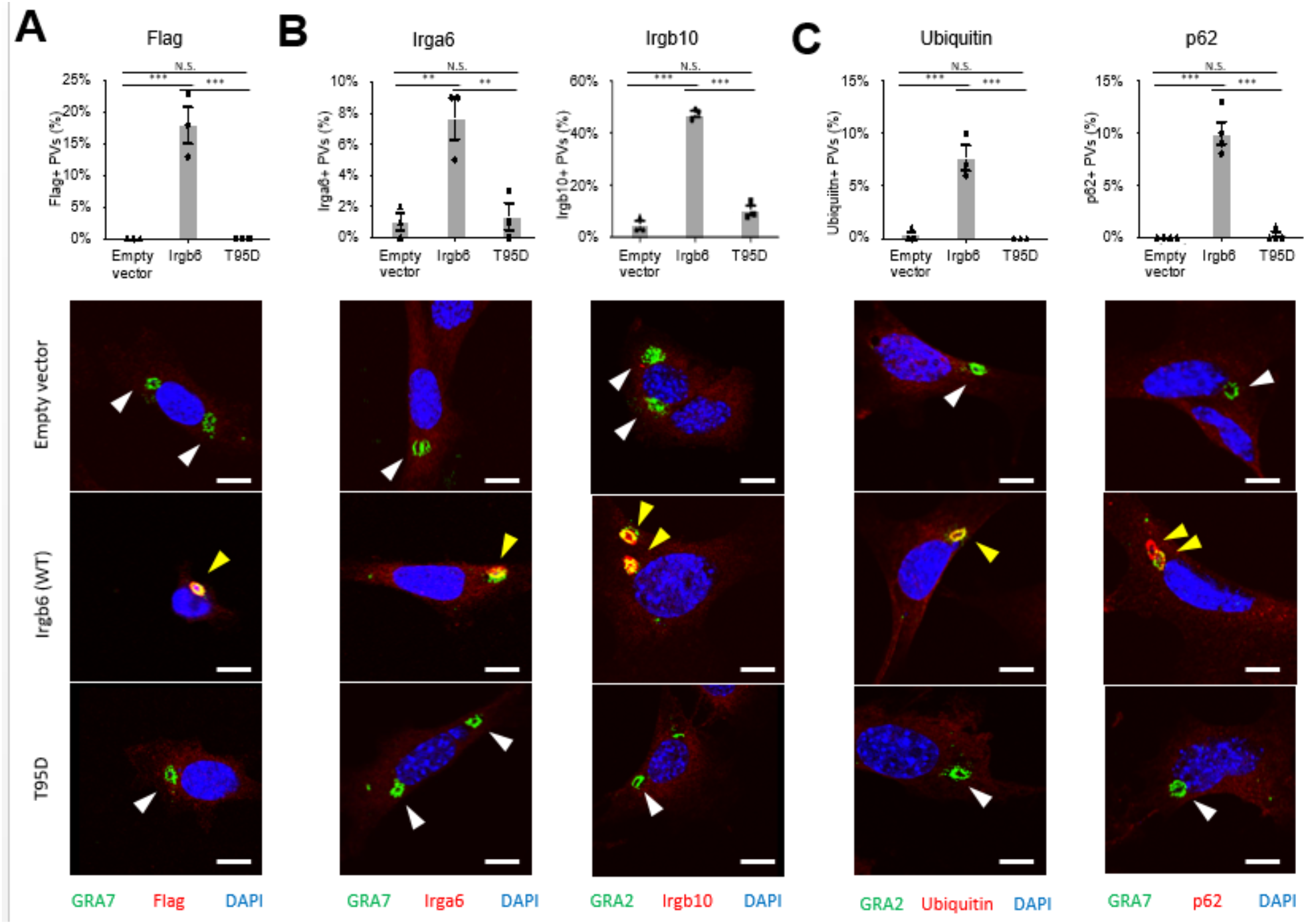
Irgb6 T95D mutation abrogates recruitment of Irgb6, other IFN-inducible GTPases and effectors. (**A** to **C**) Confocal microscope images of Flag (A) or indicated proteins (B, C) accumulation on PVs in MEFs after parasite infection. Percentage of Flag- or indicated proteins-positive PVs in cells infected with ME49 parasites. A total of 100 PVs were counted in each sample. All images are representative of three independent experiments. White arrowheads indicate PVs stained with Flag (A) or indicated proteins (B, C). Yellow arrowheads indicate *T. gondii* stained with Flag (A) or indicated proteins (B, C). Scale bars, 10 μm.

### Irgb6-T95D mutation disrupts GTPase activity

We then examined the GTPase activity of Irgb6-T95D purified through anion exchange chromatography and compared the result to that of Irgb6-WT. Size exclusion chromatography (SEC) of purified Irgb6-T95D produced two peaks, similar to Irgb6-WT (Fig 2A). SDS-PAGE analysis indicated that Irgb6-T95D was mainly eluted in the second peak, which was used for *in vitro* functional and structural assays.

**Fig 2.**
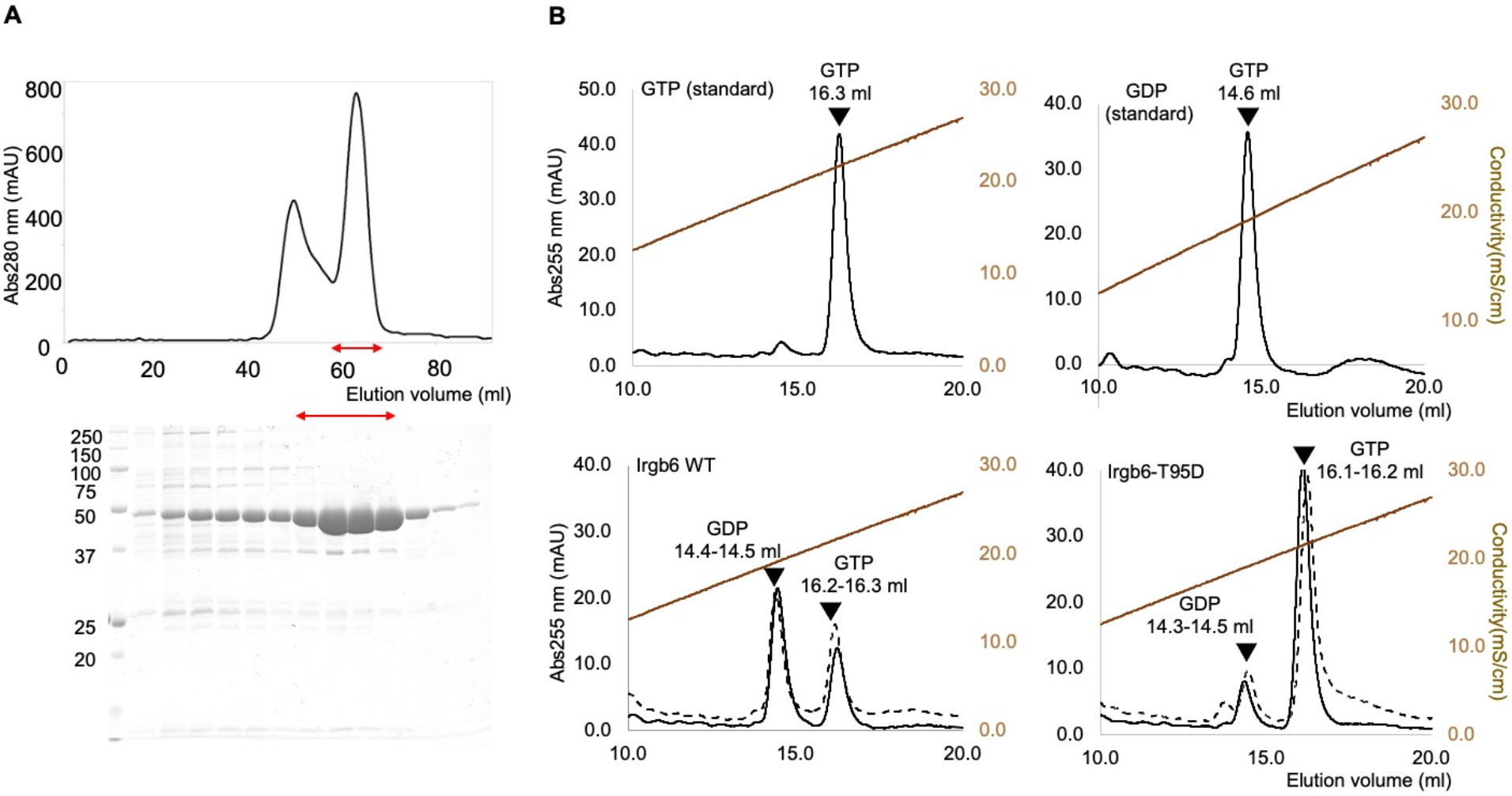
GTPase activity of Irgb6-T95D. (**A**) Purification of Irgb6-T95D. Chromatogram of SEC analysis (top) are shown with SDS-PAGE analysis (bottom). (**B**) Analyses of nucleotide components through the anion exchange chromatography. More GTP was hydrolyzed into GDP by incubating Irgb6 wild type (bottom left) than Irgb6-T95D (bottom right) at 37 °C for 30 min.

GTPase activity was determined by the extent of GTP hydrolysis to GDP after co-incubation of Irgb6 and GTP for 30 minutes. Consequently, Irgb6-WT hydrolyzed two-thirds of GTP to GDP, whereas Irgb6-T95D hydrolyzed only about 10% of GTP to GDP (Fig 2B). This indicates that the phospho-mimicking mutation of Thr95 (T95D) in the switch I region of Irgb6 significantly reduced the GTPase activity of Irgb6.

### Crystal Structures of Irgb6–T95D in the GTP-bound and nucleotide free states

The experiments above showed that phosphorylation of Thr95 in the switch I region, which was mimicked by the T95D mutation, inhibited the recruitment of Irgb6 and related molecules, including ubiquitin, to the *T. gondii* PVM. The T95D mutation also significantly reduced the GTPase activity of Irgb6, consistent with previous reports [13–14]. While the loss in GTPase activity was expected, given the location of residue 95 in the G domain, the membrane binding region in the C-domain is located more than 50 Å from residue 95 and cannot be so easily explained. In order to elucidate how the T95D mutation induced conformational changes in Irgb6 that both inhibited GTPase activity and membrane recruitment, we next performed X-ray crystallographic analyses of Irgb6-T95D in the GTP-bound and nucleotide free states.

The crystal structures of Irgb6-T95D were solved with GTP (Irgb6-T95D-GTP) and without nucleotide (nucleotide free: NF) (Irgb6-T95D-NF) at 1.68 and 2.05 Å resolution, respectively (Fig 3A and 3B and Table 1). The former structure possessed GTP in the nucleotide binding pocket (Fig 4A). The latter had neither nucleotide nor ion in the pocket (Fig 4B and 4C).

**Fig 3.**
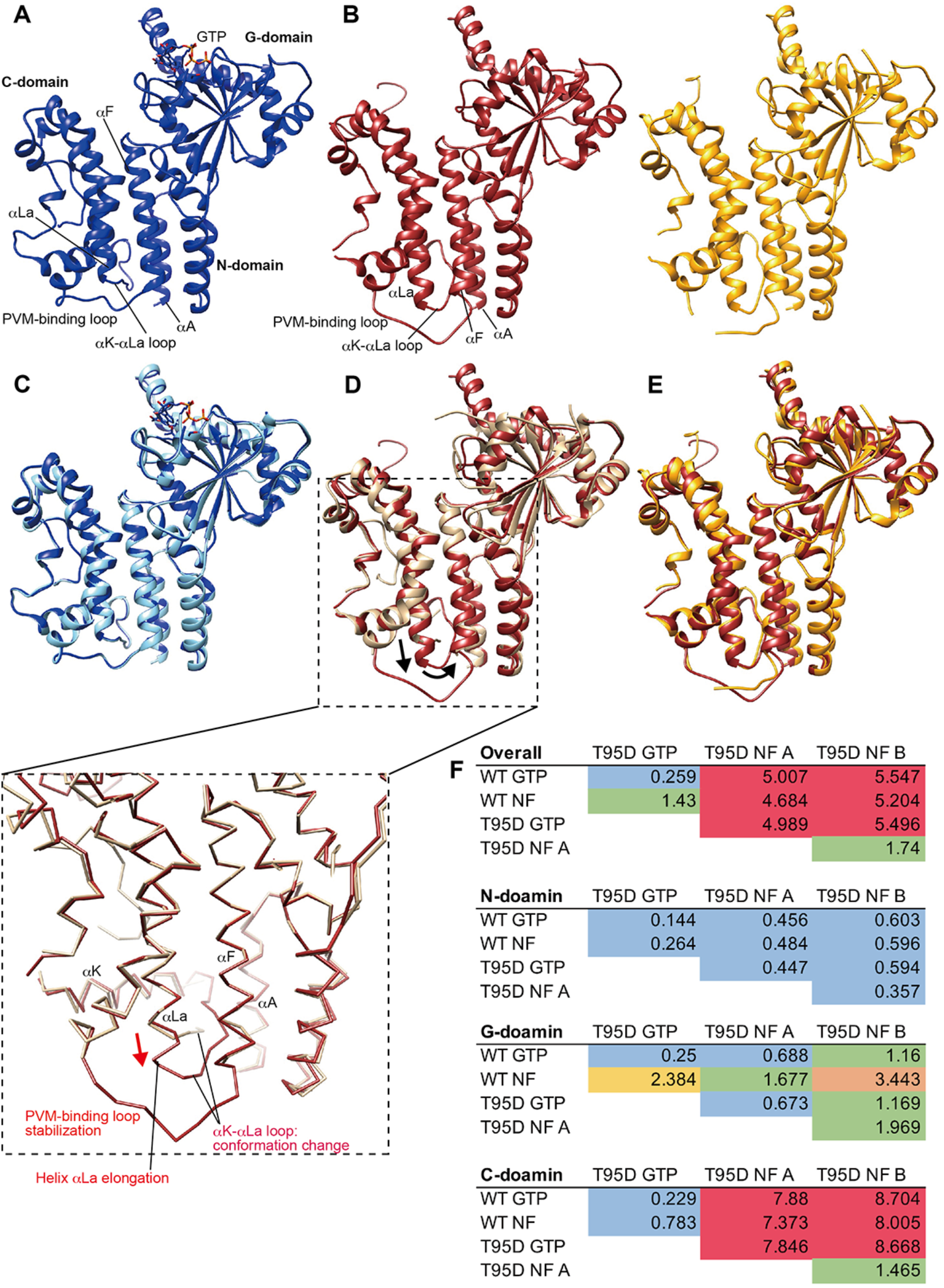
Crystal Structures of Irgb6–T95D with GTP and without any nucleotide. (**A**) Crystal structure of Irgb6-T95D-GTP. (**B**) Crystal structure of Irgb6-T95D-NF. Left, Irgb6-T95D-NF-molA; right, Irgb6-T95D-NF-molB. (**C**) Comparison of Irgb6-T95D-GTP (blue) with Irgb6-WT-GTP (sky blue). (**D**) Comparison of Irgb6-T95D-NF-molA (red) with Irgb6-WT-NF (light brown). (**E**) Comparison of Irgb6-T95D-NF-molA (red) with Irgb6-T95D-NF-molB (yellow). (**F**) RMSDs among Irgb6-WT and Irgb6-T95D. Overall RMSDs as well as RMSD values superimposed on their N-, G-, and C- domains are shown. RMSDs ≦ 1: blue, 1<RMSDs ≦ 2: green, 2<RMSDs ≦ 3: yellow, 3<RMSDs ≦ 4: orange, 4<RMSDs: red.

**Fig 4.**
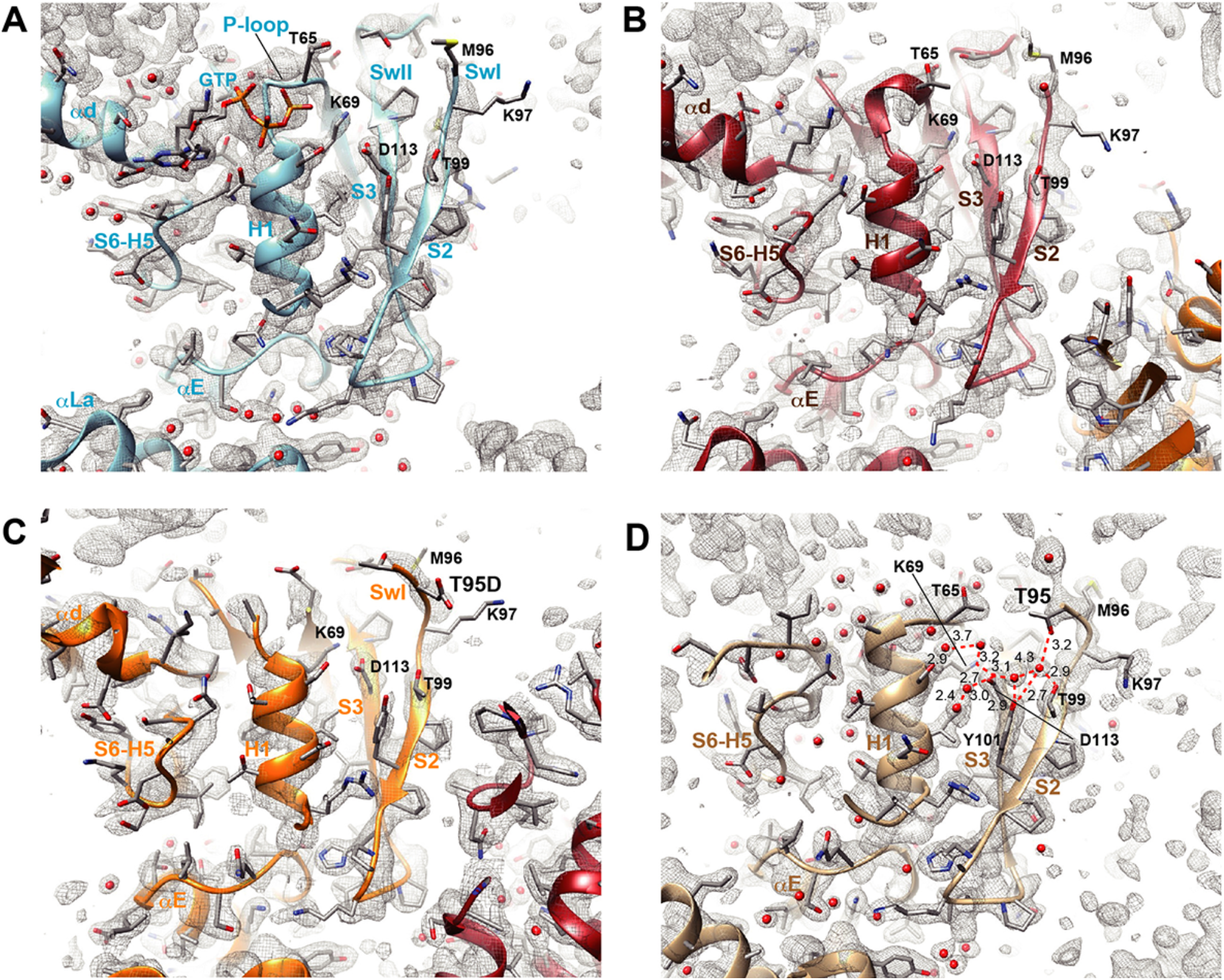
Nucleotide binding pocket of Irgb6-T95D. The 2fo-fc maps around the nucleotide binding pocket were shown at the contour level of 1.2 σ. (**A**) Irgb6-T95D-GTP. Phosphates in GTP are buried in P-loop. (**B**) Irgb6-T95D-NF-molA. P-loop takes a similar structure as Irgb6-T95D-GTP. T95D is invisible because of its flexibility. (**C**) Irgb6-T95D-NF-molB. P-loop is invisible because of its flexibility. T95D faces outward and water-mediated hydrogen bonding was not observed. (**D**) Irgb6-WT-NF. Compared to Irgb6-T95D-GTP (A), the loop-to-helix transition of the P-loop was observed. T95D faces toward the P-loop and contributes to dense water-mediated hydrogen bonding.

**Table 1.** Data Collection and Refinement Statistics.

|  | Irgb6-T95D-GTP | Irgb6-T95D-NF |
| --- | --- | --- |
| **Data collection** |  |  |
| Space group | P2<sub>1</sub>2<sub>1</sub>2<sub>1</sub> | P2<sub>1</sub>2<sub>1</sub>2<sub>1</sub> |
| Unit cell parameter |  |  |
| a, b, c (Å) | 69.122, 75.750, 80.277 | 71.532, 80.613, 147.649 |
| α, β, γ (°) | 90, 90, 90 | 90, 90, 90 |
| Resolution range (Å) | 50.000–1.67 (1.78–1.68) | 50.00–2.04(2.17–2.05) |
| Completeness (%) | 99.8 (99.1) | 99.4 (97.6) |
| R-merge(%) | 12.2 (191.6) | 11.8 (177.1) |
| R-meas (%) | 13.2 (207.2) | 12.8 (190.8) |
| <I/σ(I)> | 11.55 (0.98) | 10.25 (0.96) |
| CC<sub>1/2</sub> | 99.8 (47.5) | 99.8 (50.8) |
| Multiplicity | 6.90 (6.93) | 6.95 (7.15) |
| **Refinement** |  |  |
| Resolution range (Å) | 37.88–1.68 (1.72–1.68) | 40.55–2.05 (2.10–2.05) |
| No. Reflections | 49012 (3396) | 54351 (3645) |
| R-work/R-free | 0.2059 (0.3844) / 0.2302 (0.4215) | 0.2256 (0.3167) /0.2828 (0.3710) |
| No. atoms |  |  |
| Protein | 3286 | 6392 |
| Water | 167 | 97 |
| Ligands | 32 | – |
| RMS (bonds) (Å) | 0.008 | 0.008 |
| RMS (angles) (°) | 0.921 | 0.886 |
| Ramachandran plot (%) |  |  |
| favored | 97.24 | 97.03 |
| allowed | 2.76 | 2.97 |
| outliers | 0.00 | 0.00 |
| Rotamer outliers(%) | 0.28 | 1.00 |
| Average B-factor (Å<sup>2</sup>) | 38.45 | 56.88 |
| Protein | 38.32 | 57.02 |
| Water | 38.48 | 47.28 |
| Ligands | 52.39 | – |

Irgb6-T95D-GTP adopted the same conformation as Irgb6-WT-GTP (Fig 3A and 3C), with an overall root mean square deviation (RMSD) from Irgb6-WT-GTP of 0.259 Å (Fig 3F). As in the Irgb6-WT-GTP structure [12], Irgb6-T95D-GTP represents a collision complex with GTP in which the switch I and II regions are located away from the γ-phosphate of GTP (Fig 4A).

In contrast, the Irgb6-T95D-NF form exhibited a considerably different conformation from other forms (Fig 3B, 3D and 3E). The crystal of Irgb6-T95D-NF included two molecules in its unit cells, referred to here as molA and molB. Both molA and molB exhibited high RMSDs with respect to Irgb6-WT-NF. A domain-by-domain comparison showed that the G-domain of Irgb6-T95D-NF appeared to be more similar to Irgb6-WT-GTP than Irgb6-WT-NF. The N- domain of Irgb6-T95D-NF was relatively similar to Irgb6-WT-NF; however, the C-domain took on a considerably different conformation (Fig 3D and 3F). Specifically, the following changes occurred in the C-domain: stabilization of the PVM-binding loop, elongation of helix α-La by one turn, and a unique conformation of the αK-αLa loop, such that it is inserted into the space between the three helices αA, αF, and αLa (Fig 3D, inset).

The stabilization of the PVM-binding loop might be affected by the crystal packing environment (S2 Fig), since PVM-binding loops from molA and molB interact with each other to form the stable dimer. On the other hand, the elongation of α-La as well as the unique αK-αLa loop are likely to be influenced by the environment surrounding the three helices αA, αK, and αLa. The conformational changes of these three helices are allosterically induced by the G-domain movement, as described below. In the next section, we focus on the structural details of Irgb6-T95D-NF.

### Irgb6–T95D adapts an atypical Apo-form

As described above, the structure of the G-domain of Irgb6-T95D-NF differed significantly from that of Irgb6-WT-NF. These differences are readily visible in the active site of the G- domain with the electron density map (Fig 4). It should be noted that the resolution of both structures was approximately 2.0 Å.

In Irgb6-WT-NF, the P-loop conformation changed significantly, which further induced conformational changes around the active site, including the loop-to-helix transition of helix αd, as reported previously (Fig 4A and 4D) [12]. Notably, a dense hydrogen-bonding network via several waters connects switch I residue Thr95 and Thr99 to helix H1 to stabilize the active site conformation in the WT-NF form (Fig 4D). On the other hand, in Irgb6-T95D-NF, T95D is flexible and cannot be visualized in molA (Fig 4B) while it faces outward in molB (Fig 4C), indicating that a stable hydrogen bonding network cannot be created. Consequently, the P-loop is in a similar arrangement to the collision GTP state (molA) or becomes flexible (molB); thus, Irgb6-T95D-NF cannot take on the physiologically required nucleotide-free form.

### Thr95 phosphorylation alters the conformation of the PVM-binding region

Next, to examine how structural changes of the active site in the G-domain propagates to the N- and C-domains, Irgb6-T95D-NF and Irgb6-WT-NF were aligned with the G-domain as a reference. Consequently, we observed a clockwise rotation of N- and C-domains around the G-domain (Fig 5A). The structural change of the active site is transmitted to αE via H1 and S2-S3, and the αE movement induces a rotation of the N- and C-domain with the C-terminal side of H5 as the pivot point (Fig 5B and S1 Movie). In this process, the rotation angle of helix αA is relatively more extensive than that of the other helix movements so that the groove between αA and αF is widened, along with the flipping of Trp3 (Figs 5B and 5C and S3). Due to these changes, the αK-αLa loop can be inserted into the widened groove, further stabilizing the elongated form of helix αLa through the hydrophobic contact between Trp3 and Val364.

**Fig 5.**
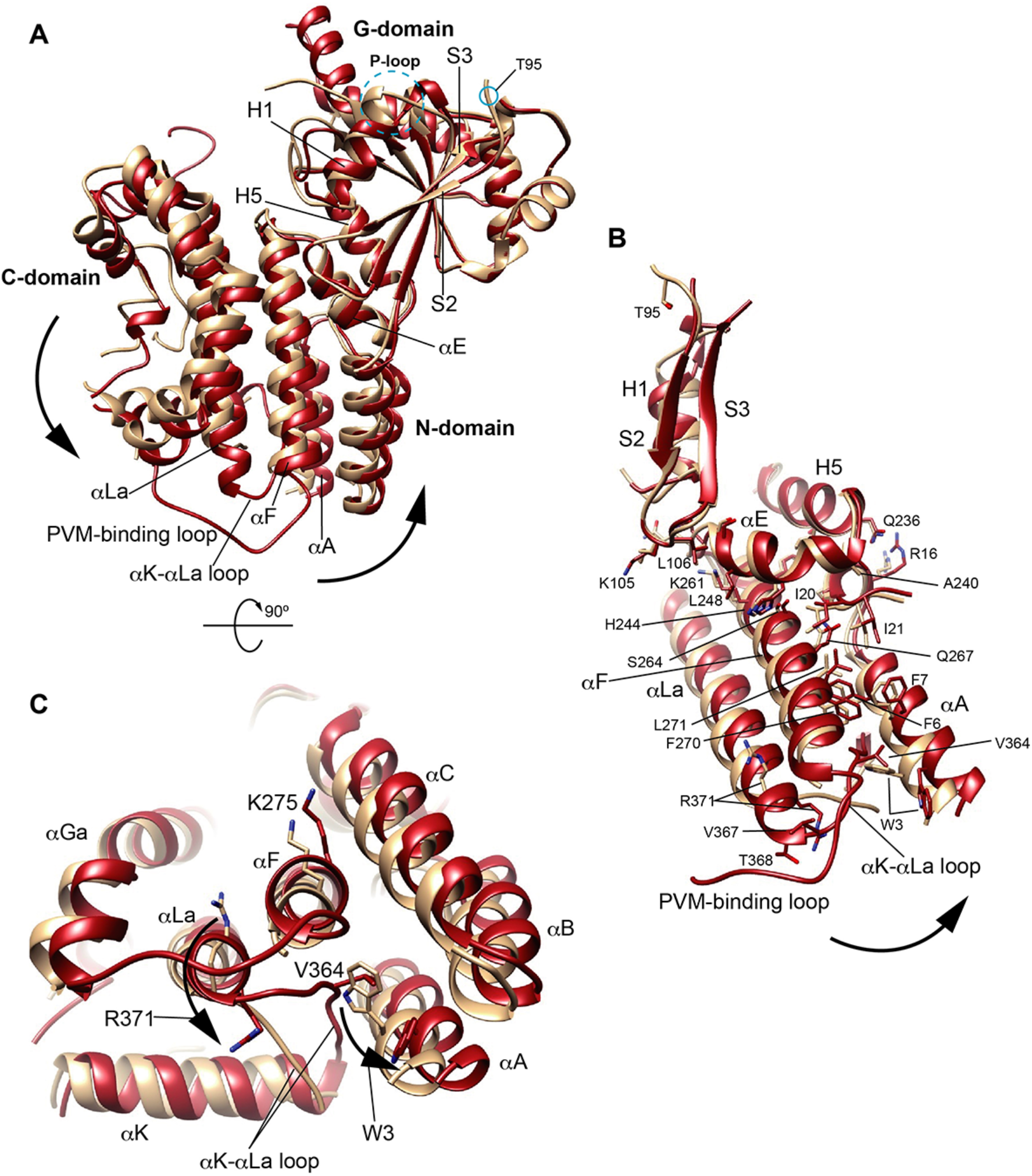
T95D-induced conformational change in N- and C- domains. (**A**) Comparison of Irgb6-T95D-NF-molA (red) with Irgb6-WT-NF (light brown). (**B**) Transduction of conformational change from the active site in the G-domain to the N- and C-domains. (**C**) Conformational change of PVM binding region. Flipping movements of Trp3 and Arg371 were observed. See Movie S1 for T95D-induced allosteric conformational change of Irgb6.

Our previous reports indicate that the amino acid triangle formed by Trp3, Lys275, and Arg371, is essential for PVM binding [11–12]. Quite suggestively, we found that Trp3 and Arg371 in Irgb6-T95D-NF have flipped and changed the structure significantly compared to Irgb6-WT-NF. Accompanying this change are significant alterations in the conformation of the PVM-binding loop as well the αK-αLa loop. These results suggest that the structure of the membrane-binding interface of Irgb6 is significantly altered by the T95D mutation.

### Phospholipid binding is drastically altered in Irgb6–T95D

Crystal structural analysis of Irgb6-T95D indicates that the T95D mutation dramatically changed the structure of the membrane-binding interface. Therefore, we examined the molecular docking of the head group of PI5P (PI5P: PubChem 643966) to the membrane-binding interface using Glide to investigate the interaction between Irgb6 and the PI5P-containing PVM (Fig 6) [17]. Since the whole structure of the PVM-binding loop was determined only in Irgb6-WT-GTP, Irgb6-T95D-GTP, and Irgb6-T95D-NF, the docking of PI5P was performed using these structures as receptors. The grid box was approximately centered on residues Trp3, Lys275, and Arg371 of Irgb6-WT-GTP. We evaluated PI5P binding with reference to the Glide scores, where a lower value indicates a more stable binding (S4 Fig). To explore the distribution of binding patterns, we extracted all poses returned by Glide using default settings.

**Fig 6.**
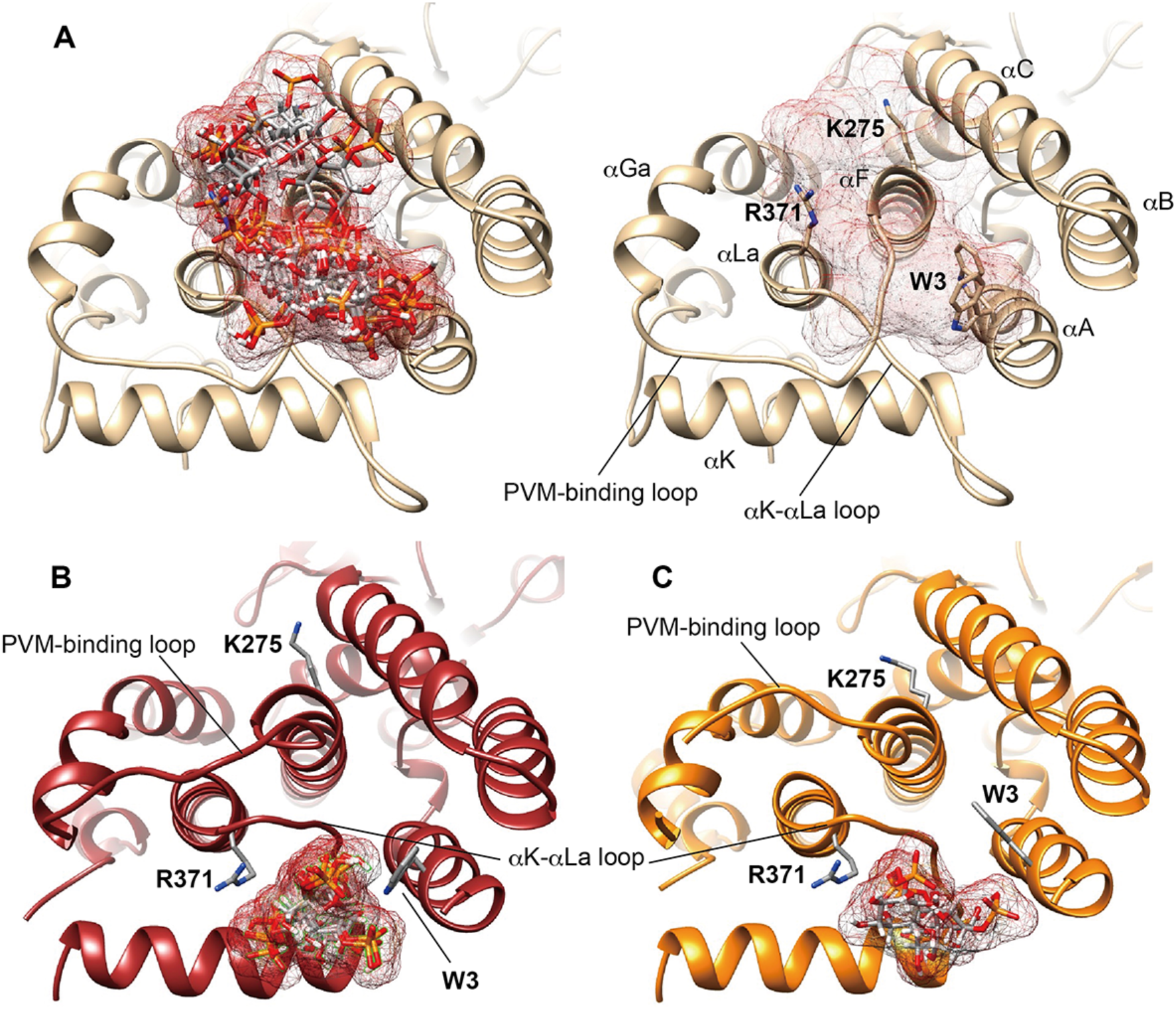
Docking simulation of PI5P to Irgb6-WT-NF and Irgb6-T95D-NF. (**A**) Docking of PI5P head to the Irgb6-WT-GTP. All poses of PI5P are overlaid (B). Molecular cloud of posing PI5P are shown in panel (C) to visualize the positions of Trp3, Lys275, and Arg371. (**B**) Docking of PI5P head to the Irgb6-T95D-NF-molA. All poses of PI5P are overlaid. (**C**) Docking of PI5P head to the Irgb6-T95D-NF-molB. All poses of PI5P are overlaid.

As previously reported, the head group of PI5P bound to the region surrounded by Trp3, Lys275, and Arg371 in Irgb6-WT-GTP. The tips of the phosphate groups of PI5P were generally oriented to bind to Arg371 or Lys 275 [Fig 6A]. The tails of the head group generally pointed to the opposite side, implying that the acyl chain would extend toward the N-terminal helices. Another characteristic of Irgb6-WT-GTP is that the PI5P binding site is relatively broad. In the case of Irgb6-T95D-NF, on the other hand, PI5P was found to bind to a completely different location outside of the original binding site (Fig 6B and 6C). This region is surrounded by R371 and Trp3, and thus this unusual pose was assumed to be due to the flipping of the side chains of these two amino acids. These results indicate that the structural changes in the membrane-bound region induced by the T95D mutation impaired normal binding of PI5P-containing vesicles, which may lead to inhibition of recruitment of Irgb6 to the PVM.

## Discussion

*T. gondii* has evolutionarily acquired a counter-defense mechanism against host cell-autonomous attacks through phosphorylation of IFN-inducible GTPases by ROP18. Our structural analysis has elucidated the atomic mechanism of how phosphorylation of Irgb6 by *T. gondii*-secreted ROP18 inactivates the GTPase activity of Irgb6. Furthermore, the structure and subsequence analysis suggest how phosphorylation induces conformational changes in the membrane-binding domain of Irgb6 at the opposite side of the protein, which, in turn, inhibit binding to the PVM.

This study found that the phosphomimetic mutation induced structural changes in the nucleotide-free state rather than to the GTP collision complex. This means that Irgb6 cannot take on the active form (which proceeds to GTP hydrolysis just after GTP binding) unless Irgb6 assumes the physiologically relevant nucleotide-free form. We recently reported that Irgb6 biochemically binds to the membrane in a nucleotide-free form and distorts the membrane through GTP binding [18]. It is consistent with the present finding that phosphorylation by ROP18 prevents Irgb6 from taking on the physiological nucleotide-free form and also changes the membrane-binding interface to inhibit membrane binding.

The conformational change in Irgb6 is transmitted to helices H5 and αE from the G-domain and further converted into rotational movement of the N- and C-domains (Fig 5). Phospholipid binding sites are located at the border between the N- and C-domains, where Lys275 and Arg371 are involved in hydrophilic binding to phospholipids, and Trp3 is involved in hydrophobic binding. The Trp3 conformation affects the magnitude of the rotation angles of the N- and C-terminal domains (S3 Fig). When the difference in rotation angle between the two domains increased by Trp3 flipping, a cleft was created between them, changing the conformation of the membrane binding interface. This is presumably a physiologically significant conformational change. *T. gondii* efficiently utilizes this strategy by phosphorylating only one threonine residue in the G-domain to open the cleft and disable the membrane binding interface.

In the molecular docking experiment, an alternative binding site for PI5P was detected because Arg371 and Trp3 were flipped in the same direction in Irgb6-T95D. Since the docking experiment was performed using the PI5P head without acyl chains, it remains to be seen what happens when the acyl chains are attached and why Irgb6-T95D could not bind to the membrane *in vivo*. In the wild-type Irgb6, PI5P-binding sites were relatively broadly distributed, whereas Irgb6-T95D has a quite narrow alternative binding site. The physiological significance of these distinct binding patterns awaits further structural analysis of membrane-bound Irgb6.

## Materials and Methods

### Immunofluorescence assay

MEFs were plated at equal densities (1.5 x 10^5^ cells per well in a 6-well plate) on glass coverslips and exposed to 10 ng/ml IFN-*γ* for 18-20 h at 37 °C. The cells were infected with ME49 tachyzoites at MOI 4 and incubated at 37 °C for 2 h. Next, cells were fixed in PBS containing 3.7% paraformaldehyde for 10 min. Cells were then permeabilized with 0.002% digitonin in PBS for 10 min and blocked with 8 % FBS in PBS for 10 min.

Cells were co-stained with anti-GRA7 and anti-Flag, Irga6, Irgb10, Ubiquitin, p62 antibodies for 1 h, followed by incubation with Alexa 488-, Alexa 594-conjugated secondary antibodies and DAPI for 30 min in the dark. Coverslips were mounted onto glass slides with PermaFluor and analyzed with confocal laser microscopy (Olympus FV3000 IX83). All procedures following 2h incubation were done in room temperature. All images were taken with a 60x objective lens.

### Mass spectrometry analysis

Irgb6-deficient MEFs stably expressing Spot-tagged Irgb6 were infected with or without *T. gondii* ME49 for 4 h and lysed in guanidine buffer (6 M guanidine-HCl, 100 mM Tris-HCl, pH 8.0, and 2 mM DTT). The lysates were diluted 8-fold with RIPA buffer (20 mM HEPES-NaOH, pH 7.5, 150 mM NaCl, 1 mM EGTA, 1 mM MgCl_2_, 0.25% sodium deoxycholate, 0.05% SDS, and 1% NP-40) supplemented with cOmplete protease inhibitor cocktail and PhosSTOP phosphatase inhibitor cocktail (Roche). After centrifugation at 20,000 × *g* for 15 min at 4 °C, the supernatants were incubated with a 5-μL slurry of anti-Spot nanobody-coupled magnetic agarose beads (Spot-Trap, ChromoTek) for 3 h at 4 °C. The beads were washed four times with RIPA buffer and then twice with 50 mM ammonium bicarbonate. Proteins on the beads were digested with 200 ng trypsin (MS grade, Thermo Fisher Scientific) at 37 °C overnight. The digests were reduced, alkylated, acidified, and desalted using GL-Tip SDB (GL Sciences). The eluates were evaporated and dissolved in 3% acetonitrile (ACN) and 0.1% trifluoroacetic acid. LC–MS/MS analysis of the resultant peptides was performed on an EASY-nLC 1200 UHPLC connected to a Q Exactive Plus mass spectrometer through a nanoelectrospray ion source (Thermo Fisher Scientific). The peptides were separated on a 75-μm inner diameter × 150 mm C18 reversed-phase column (Nikkyo Technos) with a linear 4–32% ACN gradient for 0–100 min, followed by an increase to 80% ACN for 10 min and finally held at 80% ACN for 10 min. The mass spectrometer was operated in data-dependent acquisition mode with the top 10 MS/MS method. MS1 spectra were measured with a resolution of 70,000, an automatic gain control target of 1e6, and a mass range from 350 to 1,500 *m/z*. HCD MS/MS spectra were acquired at a resolution of 17,500, an automatic gain control target of 5e4, an isolation window of 2.0 *m/z*, a maximum injection time of 60 ms, and a normalized collision energy of 27. Dynamic exclusion was set to 20 s. Raw data were directly analyzed against the SwissProt database restricted to *Mus musculus* supplemented with amino acid sequences of Spot-tagged Irgb6 and *T. gondii* ME49 proteins (ToxoDB release 38) using Proteome Discoverer v2.5 (Thermo Fisher Scientific) with the Sequest HT search engine. The search parameters were as follows: (a) trypsin as an enzyme with up to two missed cleavages; (b) precursor mass tolerance of 10 ppm; (c) fragment mass tolerance of 0.02 Da; (d) carbamidomethylation of cysteine as a fixed modification; (e) acetylation of the protein N-terminus, oxidation of methionine, and phosphorylation of serine, threonine, and tyrosine as variable modifications. Peptides were filtered at a false discovery rate of 1% using the Percolator node.

### Protein expression and purification

To create a GST-tagged Irgb6-T95D expression plasmid (pRN203), a pRN108 plasmid which is a GST-tagged Irgb6-WT expression plasmid (Saijo-Hamanio et al, 2021) was PCR-amplified with specific primers (5 ‘-<u>CCACTGGC GCAATAGAGACA</u>GAT<u>ATGAAGAGAACTCCA</u>-3’ and 5’-<u>TGTCTCTATTGCGCCAGTGG</u>-3’; the original sequence of pRN108 is underlined), and then ligated by Hi-Fi DNA assembly (New England Biolabs Inc.). Irgb6-WT and Irgb6-T95D were expressed and purified as described (Saijo-Hamano et, al. 2021) Briefly, protein expressed in *E. coli* Bl21(DE3) was purified by affinity chromatography using Glutathione Sepharose 4B column (Cytiva) and treated with GST-tagged HRV 3C protease (homemade) on the resin. The free protein was further purified by size exclusion chromatography (SEC) using a HiLoad 16/600 Superdex 75 column pg column (Cytiva).

### Analysis of nucleotide component

Nucleotide component by GTPase activity was analyzed as described (Saijo-Hamano et, al. 2021). Protein sample and GTP were prepared 200 μM in 50 mM HEPES-KOH, pH 7.5, 1 mM MgCl_2_, 1 mM EGTA-KOH, pH 7.0, and 150 mM NaCl. A 25 μl Irgb6 sample were mixed to equal volume of GTP sample and incubated at 36°C for 30 min. A 1 ml of 8 M urea was added to the mixture and heated at 95°C for 1 min, followed by ultrafiltration using Amicon Ultra-0.5 10-kD MWCO concentrator (Merck Millipore). A 900 μl of the solution that passed through the ultrafiltration membrane was analyzed by anion exchange chromatography using a Mono Q 5/50 Gl column (Cytiva) equilibrated with 50 mM HEPES-KOH, pH 7.0. Components of the reaction mixture, GTP and GDP, were completely separated by elution with 0–0.2 M NaCl gradient in 50 mM HEPES-KOH, pH 7.0. Fresh GTP (Nacalai Tesque) and GDP (WAKO) were used to confirm the elution position. A control experiment was performed using the reaction buffer.

### Crystallization

Irgb6-T95D was concentrated to 8 mg/ml in SEC solution consisting of 20 mM Tris-HCl pH7.5 (WAKO), 5 mM MgCl_2_ (WAKO), 150 mM NaCl (Nacalai Tesque), 2 mM dithiothreitol (Nacalai Tesque). Protein concentration was estimated by assuming an A280 nm of 0.916 for a 1 mg/ml solution. Nucleotide-free Irgb6-T95D crystals diffracting to 2.05 Å resolution were obtained from sitting drops with a 0.5 μl of protein solution and a 0.5 μl of reservoir solution consisting of 0.2M Sodium malonate pH7.0 (Molecular Dimensions), 20% Polyethylene Glycol 3350 (Molecular Dimensions) at 20°C. GTP-binding Irgb6-T95D crystals diffracting to 1.68 Å resolution were obtained from sitting drops with a 0.5 μl of protein solution containing 2 mM GTP (Roche) and a 0.5 μl of reservoir solution consisting of 0.2 M Sodium sulfate (Molecular Dimensions) and 20% (w/v) polyethylene glycol 3350 (Molecular Dimensions) at 20°C.

### Data collection and structure determination

Single crystals were mounted in LithoLoops (Protein Wave) with the mother liquor containing 10% (v/v) glycerol as a cryoprotectant and were frozen directly in liquid nitrogen before X-ray experiments. Diffraction data collection was performed on the BL32XU beamline at SPring-8 using the automatic data collection system ZOO (*19*). The diffraction data were processed and scaled using the automatic data processing pipeline KAMO (*20*). The structure was determined using PHENIX software suite (*21*). Initial phase was solved by molecular replacement using the Irgb6-WT crystal models (PDB ID: 7VES and 7VEX) with phenix.phaser. The initial model was automatically constructed with phenix.AutoBuild. The model was manually built with Coot (*22*) and refined with phenix.refine. The statistics of the data collection and the structure refinement are summarized in Table 1. UCSF Chimera (*23*) was used to create images and compare structures.

### In situ docking simulation

#### Protein structure preparation

The *Protein preparation wizard* from Schrödinger suite^1^ was used to prepare IRGb6-WT-GTP and Irgb6–T95D in GTP-bound and nucleotide free protein structures using default parameters. There were 2 positions for TRP3 with average occupancies of 0.53 and 0.47. Protein structures with each TRP3 position were generated. Since Incorporation of active site water molecules in the docking process is challenging^2^, protein structures with and without water were generated. In addition, protein structures with the ARG371 sidechain remodeled by Scwrl4 (PMID: 19603484) were prepared for each protein structure. Thus, for each protein, 8 structures were docked (TRPA/B x Water/NoWater x ARG371Remodeled/NotModeled). The Grid box was centered on the midpoint of TRP3, LYS275 and ARG371 using *receptor grid generation* from Schrödinger suite.^1^ The protein grids were 20 × 20 × 20 Å in size. Eight grids were generated for the 8 structures.

#### Ligand Preparation

The Pi5P molecule was truncated up to the polar head before being prepared. *LigPrep* from Schrödinger suite^3^ was used to produce low-energy, three dimensional (3D) ligands with correct chirality. Three conformations were generated and used for docking.

#### Molecular Docking

The three Pi5P head conformations were docked to the 8 grids using *Glide* (Grid-based Ligand Docking with Energetics) from the Schrödinger suite.^4^ Glide measures the ligand-receptor binding affinity in terms of Glide score. For IRGb6_T95D_nf protein, Pi5P was docked to each chain separately. In the current study, since we were focused on the global pattern of ligand distribution around the active site of the protein, and used all ligand poses, selected by Glide (usually 2-3 per docking run).

## Acknowledgments

We thank K. Chin, T. Setsu, Y. Sakihama, and T. Shimizu for assistance and other colleagues for discussions. We also thank Dr. Kohei Nishino for technical assistance in LC-MS/MS analysis. This research was partially supported by Platform Project for Supporting Drug Discovery and Life Science Research (Basis for Supporting Innovative Drug Discovery and Life Science Research (BINDS)) from AMED under Grant Number JP21am0101070.

## Funding

Japan Agency for Medical Research and Development (AMED) JP20fk0108137 (MY) Japan Agency for Medical Research and Development (AMED) JP20wm0325010 (MY)

Japan Agency for Medical Research and Development (AMED) JP20jm0210067 (MY)

Japan Agency for Medical Research and Development (AMED) JP20am0101108 (DMS)

Japan Agency for Medical Research and Development (AMED) JP21gm0810013 (RN)

Japan Society for the Promotion of Science 21K06988 (YS-H)

Japan Society for the Promotion of Science 20B304 (MY)

Japan Society for the Promotion of Science 19H04809 (MY)

Japan Society for the Promotion of Science 19H00970 (MY)

Japan Society for the Promotion of Science 21H05254 (RN)

Japan Society for the Promotion of Science 21K19352 (RN)

Japan Society for the Promotion of Science 22H02795 (RN)

Japan Science and Technology Agency (Moonshot R&D) JPMJMS2024 (RN)

## Author contributions

Conceptualization: HO, YS-H, MY, RN

Methodology: YS-H, NS, HK, DMS, MY

Investigation: HO, YS-H, AAS, EH, NS, TK, TI, EN, MS, HK, DMS

Supervision: YM, HK, DMS, MY, RN

Writing—original draft: HO, YS-H, AAS, HK, DMS, MY, RN

Writing—review & editing: HK, DMS, MY, RN

## Data and materials availability

The crystal structure data of nucleotide-free Irgb6-T95D and GTP-bound Irgb6-T95D have been registered in the Protein Data Bank (PDB) on PDBID_8H4O and PDBID_8H4M, respectively.

